# Extinction of innate floral preferences in the pollinator *Eristalis tenax*

**DOI:** 10.1101/2024.09.20.613821

**Authors:** Deepa Rajan, Aditi Mishra, Maansi Sharan, Gauri Gharpure, Shannon Olsson

**Affiliations:** University of California, San Francisco, Department of Biochemistry and Biophysics, San Francisco, California, USA; National Centre for Biological Sciences, Tata Institute of Fundamental Research, Bangalore, Karnataka, India; Uppsala University, Uppsala, Sweden; the echo network, Danish Academy of Technical Sciences, Copenhagen, Denmark

**Keywords:** Learning, multimodal, vision, olfaction, insects

## Abstract

Innate behaviors allow solitary animals to complete essential tasks in the absence of social learning. However, we know little about the degree to which ecologically relevant innate preferences can change. The hoverfly *Eristalis tenax*, a solitary generalist pollinator, is an ideal model for studying innate behavior in a naturalistic context because its survival depends on the innate ability to identify flowers. Innate behavior in *E. tenax* has previously been considered inalterable, but we hypothesized that *E. tenax* can modulate their innate behavior after training in a multimodal sensory context, in contrast to the prior work that employed unimodal sensory cues. To test this, we examined if *E. tenax* can extinguish an innate proboscis extension response (PER) to a multimodal floral object after undergoing aversive conditioning with quinine, and if flies can acquire PER to an innately unattractive object using sucrose as reinforcement. Finally, we assessed long-term memory retention. Here, we report a complete extinction of the proboscis extension response (PER) to an innately attractive floral object following aversive training. *E. tenax* can also acquire PER to an innately unattractive object after appetitive training. Flies can retain these memories for days after training, and aversive memories last longer than appetitive memories. Our results contrast with literature stating that innate preferences cannot be extinguished in *E. tenax*. This could be because our study uses multimodal objects instead of the unimodal stimuli used in previous work. Ultimately, these findings improve our understanding of how animals navigate the uncertainties of dynamic objects in the natural world.

## INTRODUCTION

Animals rely on innate behavior to find food, mates, and avoid predators in the complex natural world prior to learning (Ewert & Burghagen, 1979; Dolev & Nelson, 2014). Fishes (Dixson, Pratchett, & Munday, 2012; Gall & Mathis, 2010; Scheurer et al., 2007) and rodents (Watanabe et al., 2022) are known to identify cues from predators from birth. Furthermore, many social behaviors in vertebrates (Wei, Talwar, & Lin, 2021), birds (Diez, Cui, & MacDougall-Shackleton, 2019), and mammals (Sakata, Catalano, & Woolley, 2022) are innate, though flexible. Even the ability of humans to perceive faces arises from an interaction between innate and learned behavior (Wilkinson et al., 2014). Innate behaviors in insects are especially critical for survival because most insects are solitary with little to no parental care (Gilbert & Manica 2015) or social learning (Arenas et al., 2009; Farina, Grüter, & Díaz, 2005). However, modulation of innate preferences is also essential in a dynamic natural world. Inflexibility in innate preference can even be maladaptive, as observed in insects like jewel beetles (Hawkeswood et al., 2005), some polarotactic insects (Horváth et al. 2010), and the butterfly *Pieris virginiensis* (Augustine and Kingsolver 2018).

Nevertheless, a complete extinction of innate preferences is rarely seen. Several innate characteristics are considered largely inalterable, such as the color preference of *Xenopus* tadpoles (Hunt et al. 2020), floral color choices of *Bombus terrestris* bumblebees (Gumbert 2000), and visual attraction to tree-like objects in *Formica rufa* wood ants (Buehlmann and Graham 2022). Although there is also a growing body of literature revealing the plasticity of innate behaviors, such as the modulation of innate attraction to food by host plant gustatory cues in moths (McCormick et al., 2016) or the reduction of innate aversion to CO2 in *D. melanogaster* in the presence of fruit odor (Faucher et al., 2006), these examples feature modifications in response intensity rather than complete extinction of innate behavior itself.

Our lack of understanding of the extent to which insects can extinguish innate preferences could stem from two factors. First, most studies focus on social organisms such as honeybees, in which innate behavior and social learning are difficult to distinguish (Arenas et al., 2009; Farina, Grüter, & Díaz, 2005). Secondly, most research has also focused on the preferences of insects towards unimodal sensory cues rather than complete multimodal objects like those found in nature. Cross-modal interactions can change the valence of stimuli and alter thresholds for learning and memory retrieval (Guo & Guo, 2005). For instance, fruit flies (*D. melanogaster*) avoid small moving targets unless they are coupled with food odor (Cheng, Colbath, & Fyre, 2019; Wasserman et al., 2015).

The hoverfly *Eristalis tenax* is an ideal model for studying innate behavior in a naturalistic, multimodal context. As solitary generalist pollinators (Nordstrom et al. 2017), *E. tenax* hoverflies rely on the innate ability to identify flowers for food and survival. Furthermore, ecologically relevant multimodal floral cues that are attractive to hoverflies have been identified by sampling flowers that are attractive to hoverflies in field studies (Nordstrom et al. 2017).

PERs have been standardized to study olfactory and visual learning and innate behavior in several pollinators like bees (K. Takeda 1961; Smith and Burden 2014) and hoverflies (Neimann 2018; Lunau et al. 2018; Lunau et al. 1994). *E. tenax* hoverflies exhibit an innate Proboscis Extension Response (PER) to yellow color stimuli (K. Lunau and Wacht 1994, Neimann et al., 2018) and to appetitive sensory stimuli (Lunau et al. 2018). This behavior is triggered by photoreceptors and by gustatory receptors on the labellum and tarsi of *E. tenax* (Wacht et al. 2020; Lunau and Wacht 1994), and helps the hoverflies access pollen sources on a flower (Wacht et al. 1996). Previous research has suggested that the innate preference of *E. tenax* for yellow is largely inalterable (Lunau et al. 2018). However, these experiments focused on unimodal artificial visual stimuli, such as painted yellow dots (Lunau et al. 2018), that do not recapitulate the multimodality of natural objects from which hoverflies forage (Nordstrom et al. 2017). We hypothesized that *E. tenax* hoverflies could alter their innate behavior in a multimodal sensory context, in contrast to prior work that employed unimodal cues.

To test this hypothesis, we examined if the hoverfly *E. tenax* can alter its innate PER to a yellow floral model when trained with quinine in an aversive conditioning paradigm. We also evaluated how quickly *E. tenax* can learn novel objects as food objects using sucrose as a reinforcement. Finally, we assessed memory retention after training. Exploring the extinction of innate preferences in hoverflies can help us understand how animals navigate the uncertainties of dynamic objects in the natural world.

## METHODS AND MATERIALS

### Study Design

To assess the plasticity of innate behaviors, we identified a minimal object that elicited innate PER from untrained *E. tenax* flies. This object was a yellow flower model coupled with a floral odor blend based on the visual and olfactory features of the Sikkim positive model standardized by Nordstrom et al., 2017 (Figure 1a). The mentioned model was attractive to multiple species of hoverflies (including *E. tenax*) across continents. When assessing innate PER to this object, we used a gray disk (Figure 1b) without odor as a control because a gray disk does not encapsulate attractive features of flowers in the wild (Nordström et al., 2017). We excluded any flies that exhibited innate PER to the gray disk because these flies tended to extend their proboscis indiscriminately to random objects. In preliminary tests, we found that 17% of naive *E. tenax* hoverflies (n=163) exhibited PER to the gray disk. Once we confirmed innate PER to the yellow model with odor in the absence of PER to the gray disk without odor, we used aversive conditioning with 0.05% w/v quinine solution. We excluded flies (n = 2) that did not stop exhibiting PER after 50 trials. After aversive conditioning, we then tested retention of this aversive memory in the trained hoverflies for up to 120 hours after training.

**Fig. 1.**
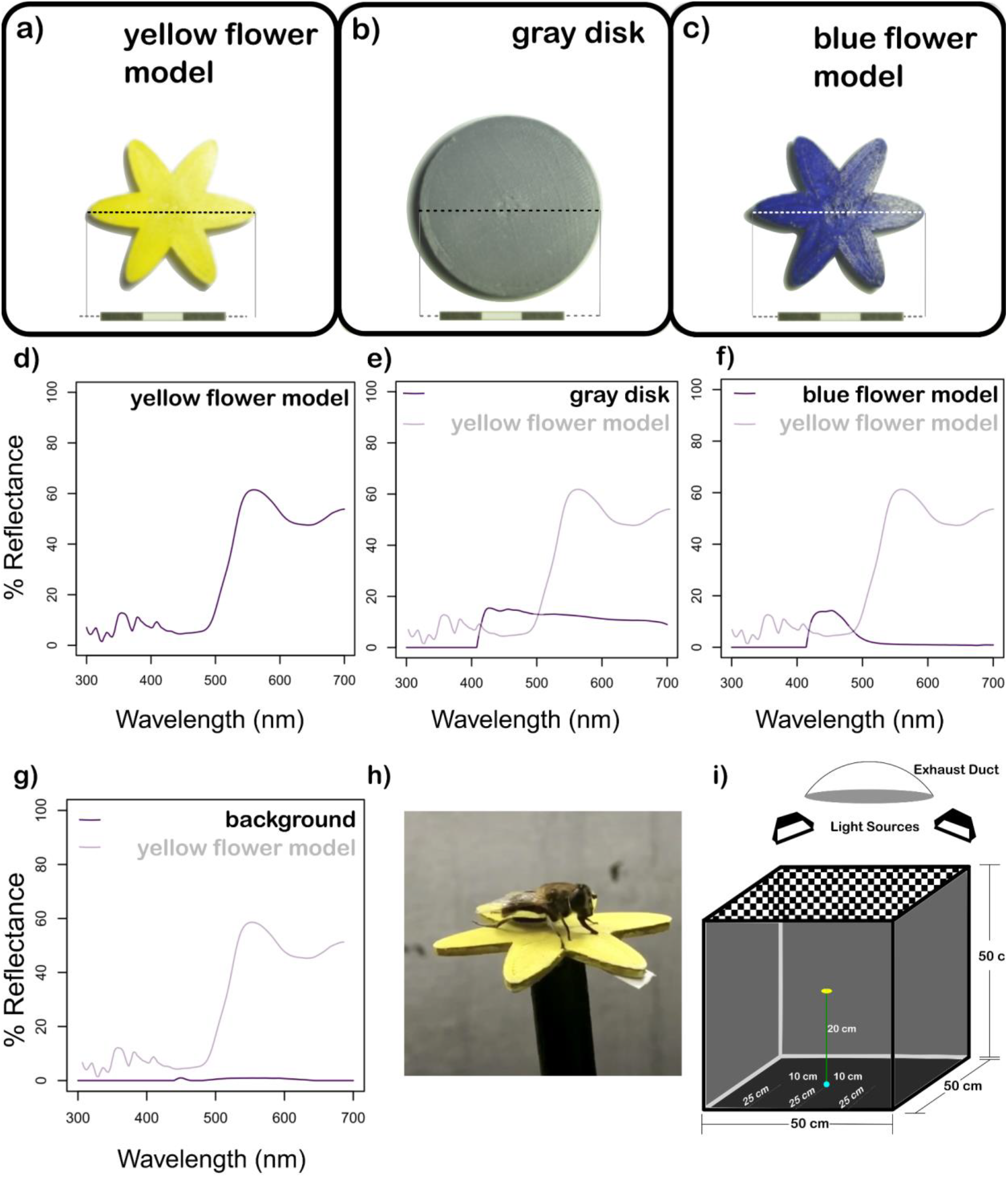
Artificial floral models used in the study. a) Innately attractive yellow flower model, b) gray disk, and c) blue flower. Scale bar = 3 cm. Reflectance spectrums of d) yellow flower, e) gray disk, f) blue flower, and g) background reflectance. h) The study organism *E. tenax* exhibiting PER to the yellow flower model with odor. i) A schematic representation of the experimental setup

After discovering that *E. tenax* can be trained to stop exhibiting PER to an innately attractive object, we checked if these flies could learn to exhibit PER to an object that is not innately attractive: a blue flower model with odor (Figure 1c). We used appetitive conditioning with 10% w/v sucrose solution. We also assessed retention of this appetitive memory for up to 120 hours after training.

### Insect maintenance

*E. tenax* pupae were obtained from Polyfly Almeria, Spain, and hatched in the lab on autoclaved vermiculite. Flies were sex-sorted upon emergence and maintained according to current literature (Nicholas et al. 2018). Up to forty hatched adult flies were housed together in white aphid-proof nylon mesh cages (32.5 cm x 32.5 cm x 32.5 cm) (BugDorm-4F3030) in a climate-controlled chamber maintained at 24 degrees Celsius, 12 hours day and night cycle, and relative humidity between 60-70%.

Adult flies were provided distilled water, 10% w/v sugar in distilled water solution, and mustard pollen mixed with powdered sugar (1:1) *ad libitum* without exposure to flowers or plants. Since hoverflies lack red-sensitive photoreceptors (An et al. 2018), the pollen-sugar mixture was mixed with carmoisine red food coloring to avoid any food-color cue associations in the hoverflies. Further, there is evidence from honeybees that while pollen can trigger PERs, it does not reinforce PER (Nicholls and Hempel de Ibarra 2013). Only flower-naïve hoverflies between the ages of 3 - 30 days following eclosion were used for experimentation. The experiments were conducted between 10 am and 7 pm during natural foraging times for these flies.

### Construction of artificial floral models

The artificial floral models used in the study were 3D printed using WoL 3D printer silk white PLA filament (WOL3D, India) and Raised N2 3D printer (Raise3D, USA) at 40% fill density to ensure that the floral models were all identical. The innately attractive yellow flower model with odor (Figure 1a) was based on a previously standardized model (Nordstrom et al. 2017) with a blend of five floral volatiles and peak reflectance in the 500 - 700 nm (Figure 1d), which appears yellow to humans. The blue floral model (Figure 1b) and the gray disk (Figure 1c) are not innately attractive to hoverflies, and have been previously standardized by Nordstrom et al. 2017. The blue flower model has peak reflectance in 400-500 nm (Figure 1f), and the gray disk model has low reflectance across 400-700 nm (Figure 1e). The 3D-printed floral models were coated with Camlin acrylic paints (KOKUYO CO., LTD.; Japan); a white base was used, followed by paints in specific ratios (Table S1) to achieve the reflectance as mentioned above.

The reflectance of all artificial floral models was recorded using a portable spectrophotometer (Ocean optics Jaz-200 spectrophotometer with an optic fiber tip of 25-cm, 600-μm Premium Fiber, UV/Visible, FL, USA). The spectrophotometer was calibrated to a white balance (Labsphere certified reflectance standard USRS-99-010 AS-01158-060 S/N 99AA08-0615-3676) under the 5000 Lux LED lights used in the experimental setup. The instrument was then used to record the hue (the band of peak wavelengths of the reflectance spectrum) and the percent reflectance of the dominant reflected wavelength of the artificial floral models used in the study (Figure 1).

All scented models have the same odor blend (Tables S2, S3). 2.5 uL of the odor solution was pipetted onto a small strip of filter paper placed underneath the surface of the 3D-printed flower so that the flies could detect the volatiles, but the filter paper strip was hidden from view. During experiments, all flower models were mounted on an artificial green stalk 20 cm high to mimic what *E. tenax* would encounter in the wild.

The volatiles emitted from the models were sampled using a 100 µm PDMS (Polydimethylsiloxane) SPME (solid-phase microextraction) fiber (model no:548575-U, SUPELCO, USA) at 5-minute intervals to check for contamination and release rate. The fiber was placed for 30 s within 10 cm of the models with the o+ volatile blend in a cleaned and heated 500 ml beaker covered with aluminum foil. The fiber was injected into the inlet of an Agilent 7890B gas chromatograph in splitless mode, and volatiles were desorbed at 270°C. An HP-5 MS column coupled with a 5977A MSD mass spectrometer (30 m × 0.25 mm i.d., 0.25 µm film thickness) was used to separate volatiles. The following temperature sequence was used: column oven 40°C for 1 min, increased to 150°C at 25°C/min, and then to 270°C at 100°C/min and held for 5 minutes. Compounds were identified by mass spectroscopy using Masshunter Qualitative Analysis software (B.07.00) and the mass spectra library NIST (National Institute of Standards and Technology and libraries created from authentic standards) (Table S3). All models used in the study were cleaned with 70% ethanol and heated at 60 degrees Celsius for 15 minutes after and before experimentation. The chemicals in the odor blend used in our study were not found in the headspace of the pollen fed to the flies (Figure S1).

### Electroantennography (EAG)

*E. tenax* between 4 - 10 days of eclosion were selected for EAG and prepared as described in Batra et al. (2019). Data was acquired using EAG2000 software, and flies were exposed to air, mineral oil solvent blank, and the odor blend pipetted onto filter paper inserted in Pasteur pipettes, as in Batra et al. (2019). A Syntech intelligent data acquisition controller (IDAC-2, SYNTECH, Netherlands) was used to collect data, and stimuli were presented as pulses of 0.5 s for 10 seconds of recording using a custom-built odor delivery system, standardized by Tait et a. 2016. The electrophysiological response of flies to the odor blend was captured in millivolts of voltage deflections. Normalized responses were calculated as the following:

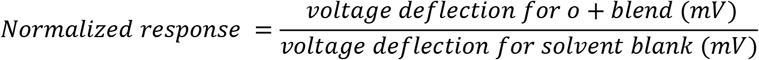

On average, the voltage deflection for the odor blend was 5.85 times higher (n = 10, sd = 2.47) than the voltage deflection for the solvent mineral oil, which indicates that the odor blend was detected at the antennae of the *E. tenax* flies since the net voltage deflection for the odor blend was significantly higher (p = 0.0011, paired t-test, n = 10, Figure S2).

### Assessing innate behavior

Flies used for aversive conditioning were starved for 24 hours to establish baseline PER motivation before introduction to the yellow model with odor. Experiments were conducted in a black Nylon mesh cage of side 50 cm (width x height x length: 50 cm x 50 cm x 50 cm) under diffused lighting from two 100 W flicker-free white LED flood lights covered with butter paper (Figure 1i). The illumination of the experimental arena was 5000 ± 50 Lux (mean ± s.e.) as measured using a lux meter. Freely moving flies were given a cotton plate with distilled water for a minute to ensure proper hydration, and then the flies were placed on the yellow flower model with odor using a wooden dowel. The flies were given up to a minute to extend their proboscis to the yellow model and the gray disk without odor (negative control). Only flies that extended their proboscis on three different placements upon the yellow model were retained for aversive conditioning. Any flies that exhibited PER to the negative control were excluded. For appetitive conditioning, flies were starved for 12 hours to ensure motivation, and then their innate behavior towards the blue model with odor was assessed using the same procedure as mentioned above. Only flies that did not show innate PER to the blue flower with odor were used for subsequent appetitive training, which constituted 85% of all the flies screened. The wooden dowel and models were cleaned using 70% ethanol and heated for 15 minutes at 60 degrees Celsius before and after experiments.

### Aversive conditioning with quinine

. Flies that exhibited innate PER to the yellow flower model with odor were then repeatedly exposed to the same yellow model along with 0.05% quinine w/v solution (2-3 droplets per petal). Flies were allowed to explore the yellow model until they touched a quinine droplet with either their legs or proboscis. One such interaction counted as a single training trial (MOV S1). Most flies exited the yellow model after touching the quinine by jumping or flying off it. After two successive training trials, the flies were tested for PER to an identical copy of the yellow model without any quinine. The flies were placed on the flower for three 1-minute testing trials, and if the flies did not exhibit PER during any of these three trials, they were considered trained. If the flies were not trained, they were provided two training trials and tested again with three testing trials. Training trials continued until the fly did not exhibit PER to the yellow flower with odor or until they completed 20 training trials, whichever happened first. Between each training trial and testing trial, the flies were given a paper towel to walk on to remove traces of quinine solution from their body. Flies that successfully completed the aversive training were given a 10% w/v sucrose solution food plate as a positive control to ensure that they could still exhibit PER to a surface that is not the yellow flower model with odor. All trained flies exhibited PER to the 10% w/v sucrose solution food plate.

### Appetitive conditioning with 10% sucrose solution

Flies were exposed to an innately unattractive model (the blue flower model with odor) with 10% w/v sucrose solution placed in droplets atop the floral model. All flies had been starved for 12 hours prior to training to establish PER motivation, and none of the flies exhibited innate PER to the models used for appetitive conditioning. Flies were allowed to explore the innately unattractive models until they touched a sucrose droplet with either their legs or proboscis, which counted as a single training trial. After two successive training trials, flies were tested for PER using an identical unrewarded model. During this testing phase, the flies were placed on the flower for three 1-minute testing trials, and the flies were considered successfully trained if they exhibited PER during all three consecutive trials. Flies were given a paper towel to walk on between the training and testing trials to remove any traces of sucrose from their body.

### Retention of aversive conditioning

Flies with innate PER to the yellow flower model with odor but not to the gray disk without odor were starved for 12 hours and subsequently given 20 successive training trials with 0.05% quinine w/v solution atop the yellow model with odor (20 trials were chosen as the training threshold for retention experiments because more than 90% of flies were trained by 20 training trials) and then were evaluated for PER on an identical yellow model without any quinine for three testing trials, each lasting one minute. Flies that did not exhibit PER during any of the testing trials were then tested for PER to the same yellow model without quinine on subsequent days. Flies that were trained at 0 hours were then tested at several time points following training: 24 hours (t1), 48 hours (t2), 72 hours (t3), 96 hours (t4), and 120 hours (t5). In between each time point, the flies experienced 12 hours of feeding and 12 hours of starvation.

During the retention testing phase, flies were placed on the yellow flower with odor for three 1-minute periods. Flies that did not exhibit PER during any of these placements were considered to have successfully retained the aversive memory. Only flies that retained aversive conditioning at any given time point were tested again in the subsequent time point.

### Retention of appetitive conditioning

We chose three trials as the training threshold for appetitive retention experiments because 50% of the flies acquired the appetitive conditioning within 2-4 training trials. Flies were given three training trials with 10% w/v sucrose on the blue flower model and then tested for three testing trials lasting one minute each for PER to an identical unrewarded blue model. Flies that exhibit PER in all three testing trials were then tested at: 24 hours (t1), 48 hours (t2), 72 hours (t3), 96 hours (t4), and 120 hours (t5). In between each time point, the flies experienced 12 hours of feeding and 12 hours of starvation. Only flies that retained appetitive conditioning at any given time point were tested again in the subsequent time point. Since flies were given 20 training trials for aversive conditioning and three training trials for appetitive conditioning, we accounted for the difference in number of exposures to the object by exposing a separate set of 32 flies to 20 training trials using 10% w/v sucrose solution placed atop the blue model with odor. These flies were tested for retention of appetitive memory using the same protocol described above.

### Statistical analysis

Exponential plateau curves were fitted using GraphPad PRISM 9 to visualize the data using the least-squares method and R^2^ values greater than 0.95. The cumulative extinction of memory between different treatments was compared using repeated measures *ANOVA* and *Fisher’s least square distance* post hoc test. Paired *t-tests* were used to statistically evaluate electroantennogram responses to the odor blend.

## RESULTS

### Innate attraction in E. tenax can be extinguished after aversive conditioning with quinine

50% of the flies that received aversive conditioning with 0.05% w/v quinine on the yellow flower model with odor stopped extending their proboscis to the model in the absence of quinine after 4 – 6 training trials, despite exhibiting innate PER to the yellow model at the beginning of the experiment. 94% of flies were trained within 20 training trials; n = 69 (Figure 2a). To account for the effects of habituation, we included a negative control experiment in which we repeated the aversive training protocol for 20 training trials without any quinine on the yellow model with odor (n = 30 flies). We also included a positive control experiment in which we repeated the aversive training protocol with 10% w/v sucrose solution atop the yellow model with odor (n = 30 trials). In both of these controls, flies did not stop exhibiting PER to the yellow flower with odor after 20 training trials.

**Fig. 2.**
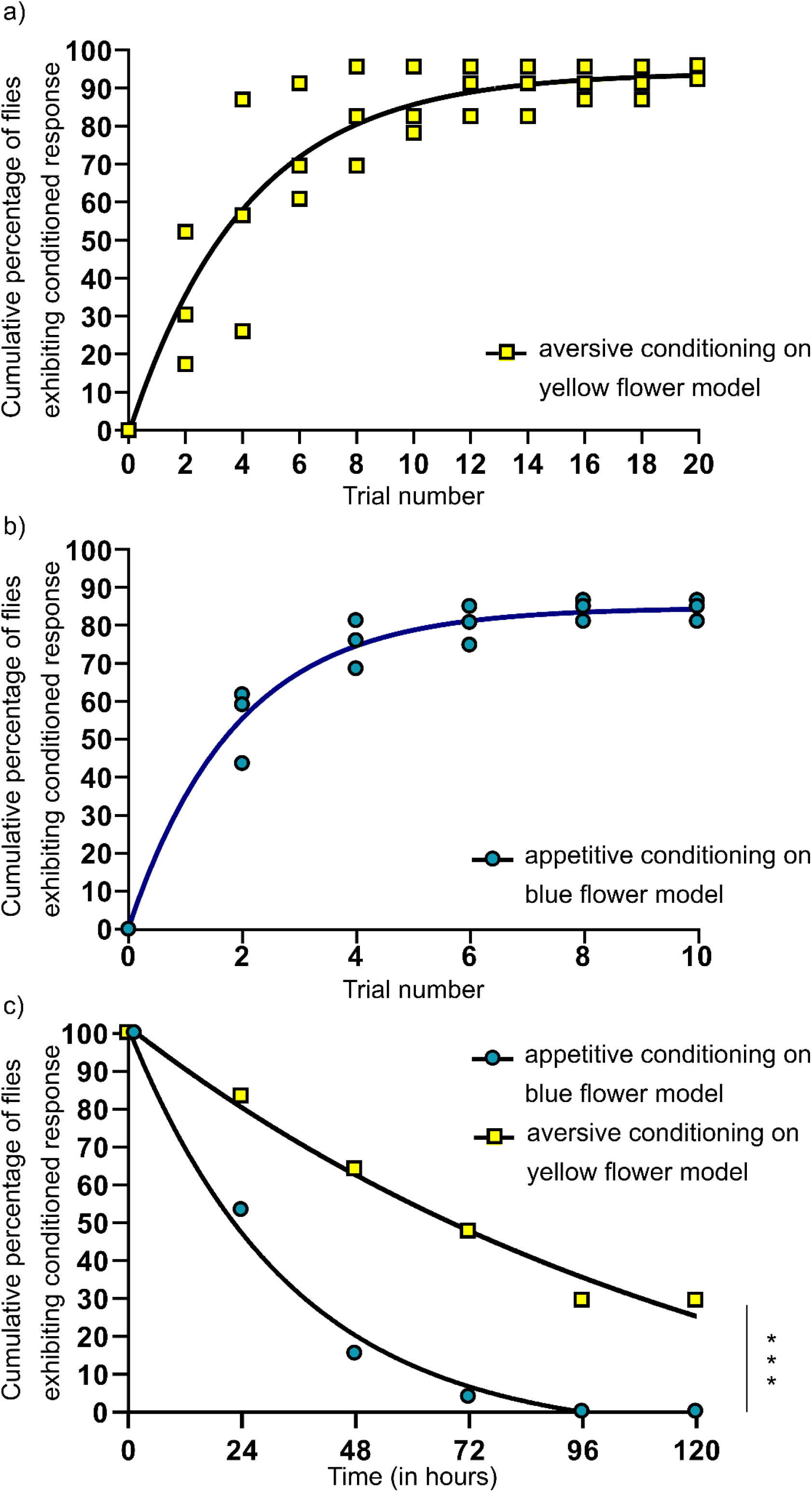
Aversive and appetitive conditioning in *Eristalis tenax*. 2a) Learning curve for aversive conditioning (3 batches containing 23 flies each; total n = 69 flies). 2b) Learning curve for appetitive conditioning (nbatch1 = 21, n_batch2_ = 27, n_batch3_ = 27 flies). 2c) Retention curves for aversive conditioning on the yellow flower model (n = 40) and appetitive conditioning on the blue flower model (n = 86) Asterisks indicate p < 0.0001, repeated measures *ANOVA*. Dotted lines show fitted curves (least-squares method, R^2^ value > 0.95)

### E. tenax flies can learn to exhibit PER to an innately unattractive object after appetitive conditioning with sucrose

50% of the flies that underwent appetitive conditioning with 10% w/v sucrose on the blue flower model with odor started extending their proboscis to the unrewarded blue model with odor within 2 – 4 training trials, despite showing no innate attraction to the model (Figure 2b). The fraction of the population that successfully learned plateaued at 85% of all flies tested (n = 75), and 11 flies did not learn to exhibit PER to the unrewarded blue model with odor.

### E. tenax flies can retain aversive training memories longer than appetitive training memories

50% of flies lost their appetitive training memory to the blue flower with odor within 24 hours (Figure 2c, Table S4, n = 86), while 50% extinction in aversive training to the yellow flower with odor only occurred at 72 hours (Figure 2c, Table S4). At 96 hours, all surviving flies showed extinction to the appetitive conditioning to the blue flower model, but 28% of flies retained aversive conditioning to the yellow flower model at the same time point (Figure 3c; Table S1). In the batch of flies that received 20 appetitive training trials on the blue flower model, 40% of the flies lost the appetitive conditioning memory by 24 hours (n = 32, Figure S3, Table S4). At 48 hours, 85% of the flies lost the appetitive training to the blue flower model, and all the surviving flies lost PER to the blue model by 96 hours (Figure S3, Table S4). The difference between the retention curves between appetitive and aversive conditioning was statistically significant (p < 0.0001, *repeated measures ANOVA*).

## DISCUSSION

Globally, hoverflies are known to visit 72% of all food crops and are robust to changes in land-use patterns and colony collapse (Doyle et al. 2020). In light of the drastic decline in insect populations (Powney et al. 2019; Rhodes 2018), hoverflies are an important asset for the ecosystem service of pollination (Rader et al. 2015). We can take a step towards mitigating the looming pollinator crisis by understanding how generalist pollinators such as hoverflies alter their innate preferences to navigate dynamic environmental challenges.

Here, we identified an extinction of innate PER to a floral model containing both visual and odor cues in 94% of *Eristalis tenax* (n = 69) after 20 aversive training trials (Figure 2a). This training was retained for at least 48 hours in more than 50% of the hoverflies (64%, n = 40, Figure 2c). With appetitive training, more than 50% of tested *E. tenax* learned to exhibit PER to an innately unattractive floral object with odor cues after only four training exposures, with ∼80% of tested *E. tenax* exhibiting PER to this object after ten training exposures (Figure 2b). Finally, the extinction of appetitive conditioning occurred significantly more quickly, with 50 % extinction at t1 = 24 hours, as compared to aversive conditioning (50% extinction at t3 = 72 hours; Figure 2c, Table S4). In particular, the difference in retention curves between aversive and appetitive memories (Figure 2c) may portend the resilience of hoverflies since they can avoid noxious chemicals for a long time while quickly taking advantage of dynamic and short-lived appetitive food sources.

Our results contrast with literature stating that innate PER to yellow colors cannot be extinguished in *E. tenax* (Lunau et al., 2018). This difference could be because of the multimodal features (visual cues in concert with odor cues) of the model used in our study as opposed to the unimodal object used in previous work (Lunau et al., 2018). Furthermore, the model used in our study was also derived from *in situ* sampling of flowers visited by hoverflies in multiple natural environments (Nordström et al., 2017) and thus incorporates ecologically relevant cues. Our work sets the stage for future research delving into how neural circuitry changes during the extinction of innate behavior in *E. tenax*. Other studies have shown that unlearning initially attractive stimuli involves the activation of aversive circuits (Jacob et al. 2021) rather than simply attenuating a reward circuit. Such neurophysiology might be apparent in *E. tenax* as well. Multisensory stimuli have also been shown to augment memory expression by binding different types of modality-specific neurons, essentially increasing the range of Kenyan cells expressing the memory engram (Okray et al. 2023). This mechanism could also apply to hoverflies, as it is consistent with our findings that multimodal sensory cues are more effective at modulating innate preferences.

In addition to playing an important role in evoking foraging behavior (Balamurali et al., 2020; Wilmsen et al., 2017; Nordström et al., 2017, Chapter 2), gustatory cues can also act as a safeguard against toxins that impair modality-specific learning (Muth, Francis and Leonard, 2019). As seen in our experiments, quinine is behaviorally relevant to pollinators, and when coupled with suitable multimodal stimuli, it can modulate innate preferences without severe physiological or learning impairment. Quinine has also been observed to deter pollen collection and cause switching in the foraging tactics of other pollinators, such as bumblebees (Muth, Francis and Leonard 2016).

Moreover, rapid appetitive learning and rapid extinction of appetitive memory, as seen in our results (Figure 2b, 2c), suggests that *E. tenax* can exploit short-lived flowers as food sources, which is consistent with the temporary floral constancies observed in some hoverflies (Goulson and Wright 1998). The frugal innate search template of hoverflies, coupled with their ability to unlearn the innate preference for the yellow floral model, could help them avoid sub-optimal, maladaptive, or noxious choices, thus making them resilient generalist pollinators. This flexibility of innate behavior gives hoverflies an advantage over other pollinators, such as honey bees, which continue to forage from unpalatable, quinine-laced flowers regardless of their energy budget (Desmedt et al. 2016).

In this vein, aversive learning via unpalatable reinforcement and modulation of innate preferences could be a robust safeguard against suboptimal or toxic food objects in many pollinators. For example, several pesticides, such as neonicotinoids, are tasteless (Raine and Gill, 2015) and undetectable except at very high concentrations (Nagloo, Rigosi, and O’Carroll 2023; Parkinson et al., 2023). Pollinators can, therefore, potentially consume large quantities of such toxins (Kessler et al., 2015; Arce et al., 2018; Nagloo, Rigosi, and O’Carroll 2023). These pesticides often contaminate pollen and nectar (Zhang et al., 2023), and are detrimental even at sub-lethal doses to pollinators (Wu-Smart and Spivak; 2016; Singla et al., 2020; Siviter, Richman, and Muth, 2021). Coupling bitter compounds like quinine with tasteless contact-poison pesticides like neonicotinoids (Anadón et al., 2020) could be a viable strategy to reduce the exposure of pollinators to such pesticides and should be explored further. This approach holds promise not only because quinine can be detected by bees (Wright et al., 2010) and flies (Sellier, Reeb and Marion-Poll, 2011) at concentrations much lower than the lethal dose, but also because the lethal dose itself for quinine (LD50 = 10mM quinine at 24 hr for bees; Wright et al., 2010) is much higher than that of neonicotinoid pesticides like clothianidin (LD50 = 0.53 ppm or 25.4 ng clothianidin per bee; Yao, Zhu and Adamczyk, 2018). The slow extinction of aversive conditioning (Figure 2c) is promising and suggests that aversive training could be long-lasting in protecting hoverflies against pesticides in an agricultural context.

Future experiments can move beyond our minimal flower model and assess the effects of more complex sensory cues on learning. Conducting these experiments in the field using the expansive range of sensory cues from natural flowers may offer a fuller picture of the degree to which innate preferences can change in other solitary species. Countering such well-worn adages as “a leopard can’t change its spots” and “a tiger can’t change its stripes,” we show here that an animal can extinguish something as fundamental as the food preference with which it was born. Moreover, a multimodal sensory context may facilitate this extinction of innate behavior. Ultimately, these findings improve our understanding of how animals navigate uncertainties in natural environments that are constantly changing, and pave the way for uncovering common principles about the mutability of innate preferences to multimodal objects.

## Acknowledgments

We thank Ashwin Suranarayan and Veena S Gowda for preliminary experiments that led to the design of this study. We would like to thank Roopa Rajendran for helping us with the maintenance and upkeep of the hoverflies. This work was supported by a Fulbright Fellowship to D.R., Stiftelsen Olle Engkvist Byggmästare, Sweden (grant numbers 2014/254, 2016/348) to S.B.O.; and NCBS-TIFR funding, Department of Atomic Energy, Government of India to S.B.O. under project no. 12-R&D-TFR-5.04-0800 and 12-R&D-TFR-5.04-0900.

## Author Contributions

S.B.O., D.R., and A.M. contributed to the study conception, design, and writing. D.R., A.M., M.S, and G.G. contributed to material preparation, data collection, and analysis.

## Data availability

Data is provided within the manuscript or supplementary information files.

