## SUPPLEMENTARY INFORMATION for "Extinction of innate floral preferences in the pollinator *Eristalis tenax*"

Journal of Chemical Ecology

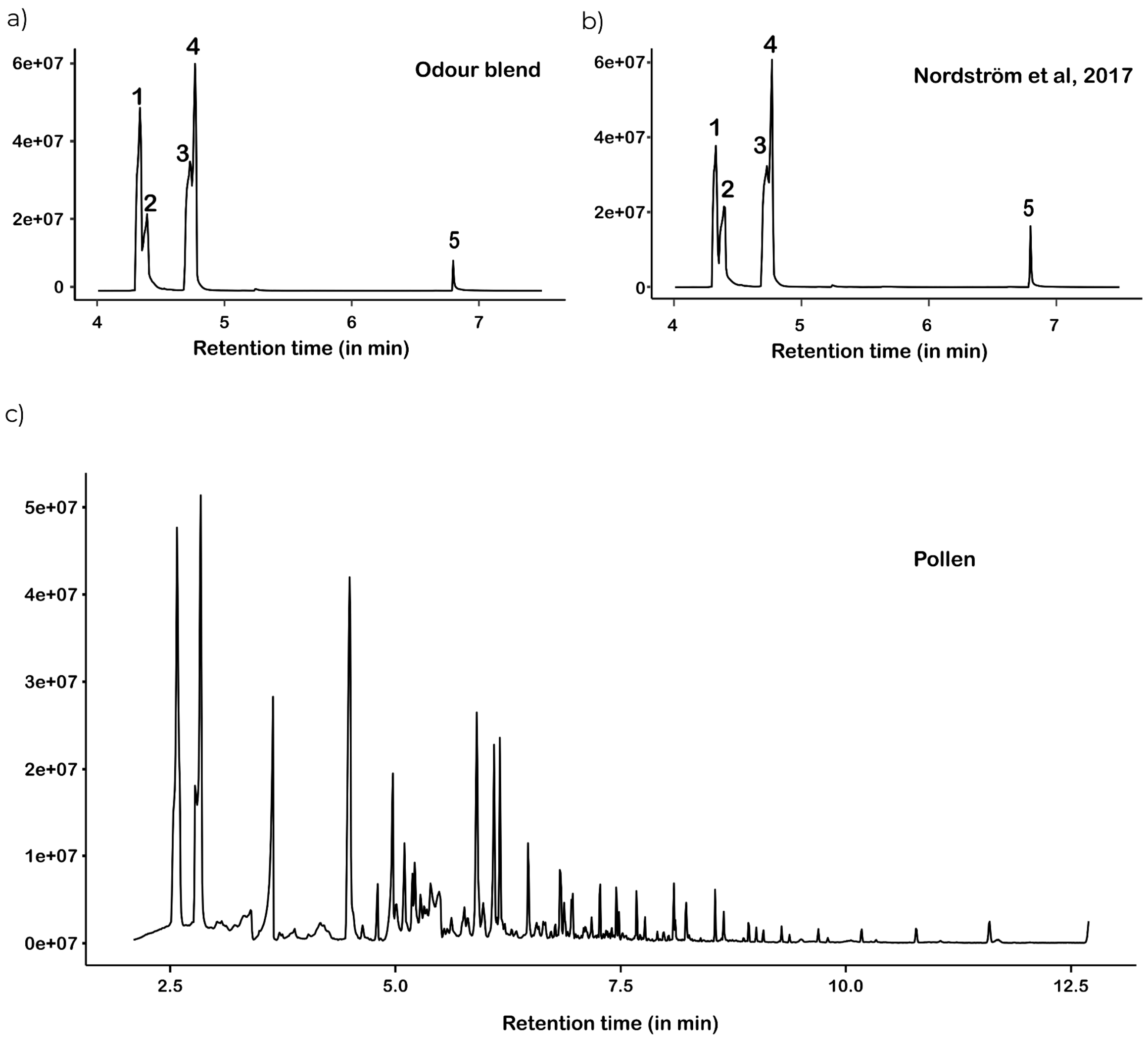


**Fig. S1** Count versus gas chromatography retention index graphs for a) the odor blends used in the study, b) previously standardized blend in Nordstrom et al. 2017, and c) the headspace of pollen samples (gas chromatography / mass spectrometry raw data traces uploaded separately)

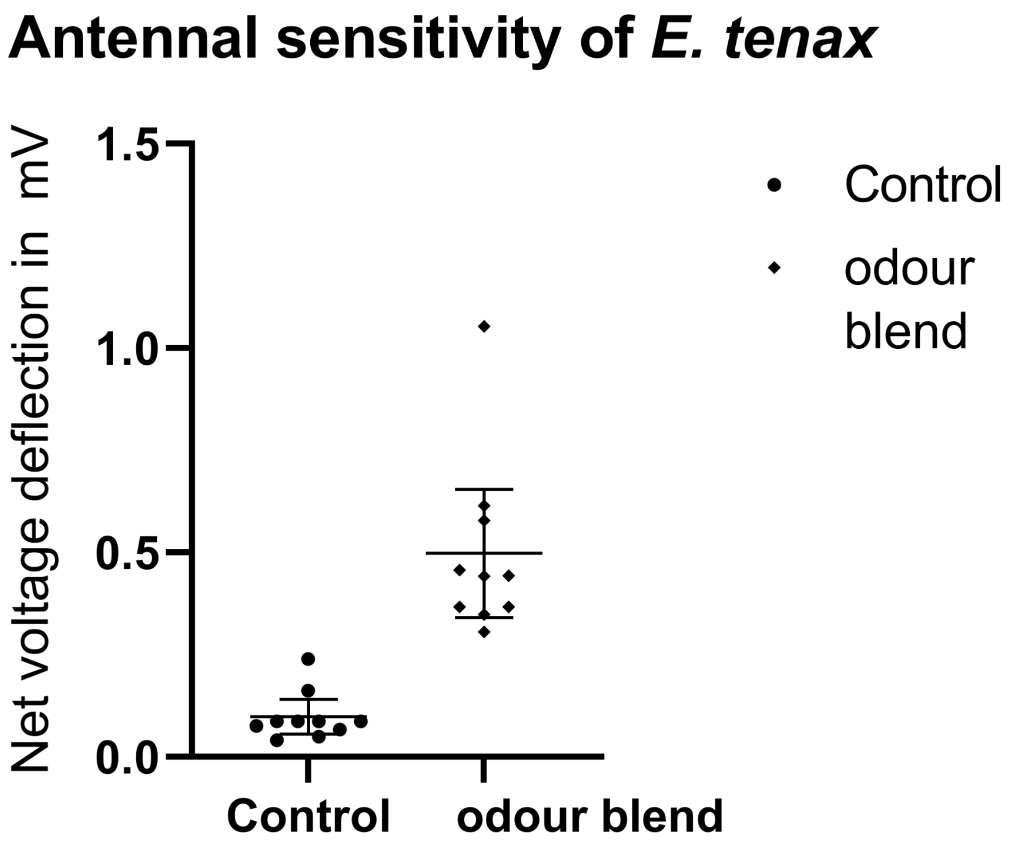


**Fig. S2** Electroantennogram of E. tenax comparing net voltage deflection for mineral oil solvent (control) and the odor blend (n = 10)

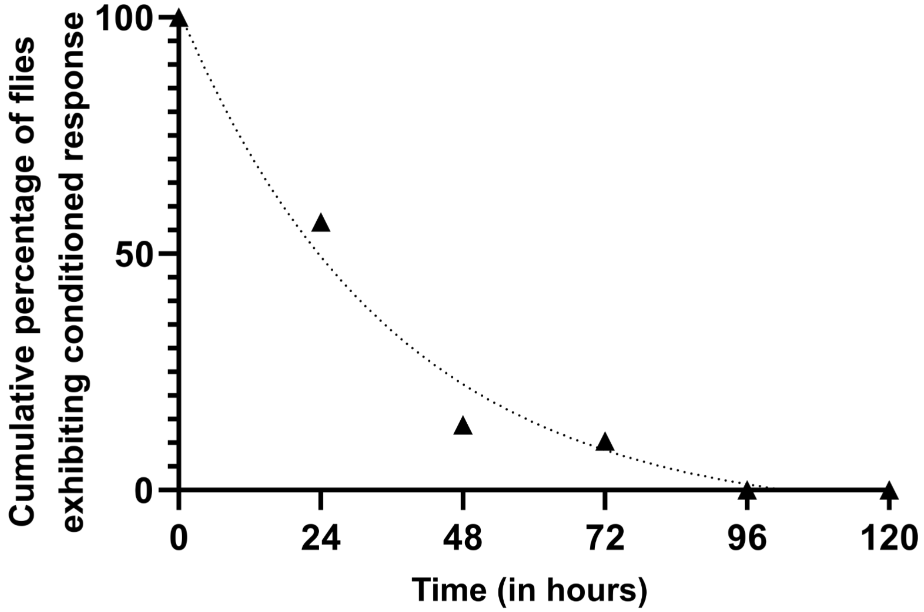


**Fig. S3** Retention of appetitive training to the blue flower model with odor after 20 training trials with 10% w/v sucrose (n = 32)

**Table S1** Ratios of paints used in each floral model

| Floral model | Blend of Camlin acrylic colors |
|---|---|
| yellow flower model with odor (reflectance peak 540-580 nm) | 14:1:3 ratio for Chrome yellow: sap green: white acrylic paint. |
| blue flower model (reflectance peak 400-440 nm) | 100% Ultramarine blue acrylic paint. |
| gray disk (reflectance peak 400-800nm, reduced spectral intensity, i.e., brightness) | 9:1 ratio of White: black acrylic paint. |

**Table S2** Composition of the volatiles used

| Odor blend | Composition |
|---|---|
| Odor blend used for learning experiments | 16.1 µl 2-Ethyltoluene [Sigma Aldrich, 99% purity]: 11.8 µl p-Cymene [Sigma Aldrich, 99% purity]: 60 µl Undecanal [Sigma Aldrich, 98% purity]: 24.1 µl r-Limonene [Sigma Aldrich, 97% purity]: 17.3 µl 6-Methyl-5-hepten-2-one [Fluka]: 82.7 µl mineral oil [Sigma Aldrich] |

**Table S3** Relative ratios of chemicals in the headspace of odor blend used in this study compared to blend from Nordström et al, 2017, which was constituted from empirically measured release rates from flowers in nature previously studied in the literature

| Blend | Retention Time | Area | Base peak | chemical | Relative ratio of areas under different chemicals of the blend |
|---|---|---|---|---|---|
| **odor blend used for learning experiments** | 4.33 | 101406628 | 105.1 | 2-Ethyltoluene | 27.59: 16.59: 53.16: 2.65 |
| | 4.39 | 60991256 | 108.1 | 6-Methyl-5-hepten-2-one | |
| | 4.77 | 195407267 | 93.1 | Limonene - Cymene | |
| | 6.8 | 9752319 | 57.1 | Undecanal | |
| **Blend from Nordström et al, 2017** | 4.32 | 77557758 | 105.1 | 2-Ethyltoluene | 22.21: 19.49: 53.48: 4.82 |
| | 4.39 | 68037452 | 108.1 | 6-Methyl-5-hepten-2-one | |
| | 4.76 | 186748875 | 93.1 | Limonene - Cymene | |
| | 6.8 | 16831435 | 57.1 | Undecanal | |

**Table S4** Retention of different training regimes (aversive conditioning: 0.05% quinine on the yellow model (20 trials); appetitive conditioning: 10% sucrose on the blue model (3 trials and 20 trials). Numbers within brackets represent the numbers in each of the two sets of flies used to study retention of appetitive training

Testn = total number of flies tested at time n,

CONn = flies that retained conditioning (i.e., flies that did not exhibit PER to yellow model at time tn or flies that exhibited PER to blue model at time tn),

EXTn = flies with extinction of conditioning, i.e., flies that exhibited PER to yellow model in aversive conditioning or flies that did not exhibit PER to blue model in appetitive conditioning at time tn,

DEADn = number of flies that died between time tn-1 and tn

Testn = Testn-1 – EXTn-1 – DEADn

| Retention of aversive training with 20 training trials (0.05% quinine on yellow model) | | | | | | |
|---|---|---|---|---|---|---|
| Time point (in hrs) | Flies that did not exhibit PER to yellow model | Flies that exhibited PER to yellow model | Number of flies dead at time point | Total number of flies tested | Cumulative number of flies with extinction of aversive training | Cumulative percentage of flies exhibiting learned behavior |
| tn | CONn | EXTn | DEADn | TESTn | CUM.EXTn | 1- $CUM.EXTn/CUM.EXTn + CO$[illegible] |
| t0 = 0 | 40 | 0 | 0 | 40 | 0 | 100 |
| t1 = 24 | 30 | 6 | 4 | 36 | 6 | 83.33 |
| t2 = 48 | 16 | 3 | 11 | 19 | 9 | 64 |
| t3 = 72 | 10 | 2 | 4 | 12 | 11 | 43.48 |
| t4 = 96 | 5 | 1 | 4 | 6 | 12 | 27.78 |
| t5= 120 | 5 | 0 | 0 | 5 | 13 | 27.78 |
| Retention of appetitive training with three training trials (10% sucrose on blue model) | | | | | | |
| Time point (hrs) | Flies that exhibited PER to blue model | Flies that did not exhibit PER to blue model | Number of flies dead at time point | Total number of flies tested at time point | Cumulative number of flies with extinction of appetitive training | Cumulative percentage of flies exhibiting learned behavior at the time point |
| tn | CONn | EXTn | DEADn | TESTn | CUM.EXTn | $CONn/ CUM.EXTn + DEADn$ |
| t0 = 0 | 86 | 0 | 0 | 86 | 0 | 100 |

| | | | | | | |
|---|---|---|---|---|---|---|
| t1 = 24 | 45 | 41 | 0 | 86 | 41 | 51.67 |
| t2 = 48 | 12 | 21 | 12 | 33 | 62 | 15.34 |
| t3 = 72 | 4 | 8 | 0 | 12 | 70 | 4.256 |
| t4 = 96 | 0 | 1 | 3 | 1 | 71 | 0 |
| Retention of appetitive training with 20 training trials (10% sucrose on blue model) | | | | | | |
| Time point (hrs) | Flies that exhibited PER to blue model | Flies that did not exhibit PER to blue model | Number of flies dead at time point | Total number of flies tested at time point | Cumulative number of flies with extinction of appetitive training | Cumulative percentage of flies exhibiting learned behavior at the time point |
| tn | CONn | EXTn | DEADn | TESTn | CUM.EXTn | $CONn/\ CUM.EXTn\ +\ DEADn$ |
| t0 = 0 | 32 | 0 | 0 | 32 | 0 | 100.00 |
| t1 = 24 | 17 | 13 | 2 | 30 | 13 | 56.67 |
| t2 = 48 | 4 | 12 | 1 | 16 | 25 | 13.79 |
| t3 = 72 | 3 | 1 | 0 | 4 | 26 | 11.54 |
| t4 = 96 | 0 | 2 | 1 | 2 | 100 | 0.00 |

**Mov. S1** Examples of aversive training trials. The hoverfly is exposed to droplets of 0.05% quinine solution on top of the yellow flower model, and subsequently flies away. (Movie uploaded separately.)
